# Orientation tests and long-term movement phenology establish the red admiral *Vanessa atalanta* as an applicable model for navigation research in migratory butterflies

**DOI:** 10.1101/2023.09.09.554419

**Authors:** Alexander Pakhomov, Roberts Jansons, Nazar Shapoval, Fyodor Cellarius, Anatoly Shapoval, Oliver Lindecke

**Author notes:** Correspondence: Alexander Pakhomov, Oliver Lindecke.

## Abstract

Animal migrations are disappearing globally, while insect populations are on alarming declines. Both ecosystem degradations, influenced by unpredictable impacts of climate change, are also exacerbated by human activities such as intensified land use and various forms of environmental pollution. Butterfly migrations may serve as sensitive indicator phenomena of these broader environmental changes. While the transcontinental journeys of one of the most famous Lepidopteran species, the North American monarch butterfly, *Danaus plexippus*, are documented in depth, they are a geographically restricted phenomenon. Comprehensive studies from other areas and on other migratory butterflies like the European red admiral, *Vanessa atalanta*, are notably sparse. In addition, the details of their navigational capacities and how they might be affected by the aforementioned changes remain largely enigmatic. Against this backdrop, we seek to establish the red admiral as a model for insect movement phenology and navigation behaviour which both might be impacted by environmental changes. Employing a combination of orientation tests, utilizing flight-simulators and free-flight trials during late summer, together with a 23-year study on movement phenology at a coastal migration flyway, the Baltic Sea coast, we offer broad insights into red admiral migration. In our experiments, butterflies exhibited a southwestern orientation on the Courish Spit and chose a south-southeastern trajectory in free-flight trials after translocation at the Latvian Baltic Sea coast. Directional records from decades-long trapping data, based on more than 16,000 individuals, match these findings. Nevertheless, we also found reverse movements to occur under some circumstances. At the same time, the observed estimated median dates of red admiral passages did change by one day only between decades, however, generally more butterflies were recorded in recent years. Our data thus suggest a certain degree of adaptability in the butterflies’ movement behaviour, indicating an innate migration schedule, possibly supported by a flexible navigational capacity. As the world is facing biodiversity loss at a high rate, long-term monitorings of indicator species become important. By establishing the red admirals as a model for butterfly migration, we expect insights into broader movement patterns and navigational strategies in Lepidoptera negotiating human-dominated environments, filling a crucial gap in our current understanding of these interdependent aspects of insect biology.

## Introduction

Every year billions of animals from a multitude of taxa, from insects to birds and mammals, perform migrations of hundreds or thousands of kilometres. Migration is a process that allows animals to escape inimical environmental conditions at their breeding grounds and to reach more favourable sites for the duration needed until returning becomes beneficial. These mass movements of taxa are characterised by precisely coordinated orientation and timing schedules that allow these animals to successfully navigate to their goal-areas (Merlin and Liedvogel, 2019). Yet, such mechanisms of orientation and navigation have been studied in-depth only in a few selected species. For example, a behavioural paradigm that allows measuring orientation in birds under controlled laboratory conditions – round cages (Kramer, 1949) and Emlen funnels (Emlen and Emlen, 1966) – has led to a better understanding of migratory bird navigation mechanisms in all parts of the world based on seasonally expressed *Zugunruhe* (Wiltschko and Wiltschko, 2015). Similar behavioural paradigms designed for other migratory vertebrates have been developed only more recently (e.g., for sea turtles (Lohmann, 1991), fish (Lohmann, 1991; Bottesch et al., 2016), and bats (Lindecke et al., 2019)). However, long-range seasonal migration is a widespread phenomenon among insects on all continents, especially in Lepidoptera (Chowdhury et al., 2021). Yet, until the beginning of the 21st century, there was also no adequate laboratory setup similar to the Emlen funnel that could help researchers to infer migratory behaviour and measure orientation of butterflies and other insects. The development of laboratory-based flight systems, such as the Mouritsen-Frost flight simulator (Mouritsen and Frost, 2002; Dreyer et al., 2021), changed that.

Most studies dealing with orientation and navigation abilities of migratory butterflies were performed on an iconic model species, the North American monarch butterfly *Danaus plexippus* (Linnaeus, 1758). During autumn, monarchs undertake a remarkably long migratory journey from their northern range in Canada and USA to overwinter in central Mexico (Urquhart and Urquhart, 1978). However, their overwintering behaviour is unique among butterflies: thousands of individuals cluster together on trees at specific sites until spring arrives. This population-wide, goal-orientated behaviour is one of the key factors that differentiates monarchs from other diurnal migratory butterflies, which are less confined in their site choices. These either overwinter in a wider geographic range in the adult, egg, or larval stage, or they die off in the colder regions with the next generation returning only from warmer areas in the spring. It is now understood that migratory monarchs mostly rely on information from the Sun using an antenna-based time-dependent sun compass system to navigate during this long journey (Perez et al., 1997; Reppert et al., 2004; Stalleicken et al., 2005; Merlin et al., 2009). Additionally, they can use the geomagnetic field as a backup compass system (Reppert and de Roode, 2018) to orient in a southward direction. It has been reported that their magnetic compass is based on the determination of the inclination angle of the Earth’s magnetic field (inclination compass) and requires UV-A/blue lights with wavelengths between 380 and 420 nm (Guerra et al., 2014). However, there is no confirmation of these findings in other independent studies: monarch butterflies were disoriented in simulated overcast conditions under natural daylight in a flight simulator (Mouritsen and Frost, 2002; Stalleicken et al., 2005) but flew in southward direction on cloudy days in the wild (Schmidt-Koenig, 1979).

Despite the fact that some Palaearctic butterflies evolved the longest regularly undertaken insect migration circuit (Hu et al., 2021), we know much less about their navigation abilities compared to American monarchs. Every year, for example, painted lady butterflies *Vanessa cardui* (Linnaeus, 1758) perform their incredible, multigenerational round-trip of about 12,000 to 14,000 km between tropical West Africa and Scandinavia (Stefanescu et al., 2016; Hu et al., 2021; Talavera et al., 2023) which strongly suggests that well-developed compass systems are at play. However, there is no direct experimental evidence that this diurnal migrant uses a time-compensated sun compass for orientation as monarchs can do (Nesbit et al., 2009): laboratory-reared painted lady butterflies were unable to maintain their southward direction under simulated overcast conditions, unlike their conspecifics from the control group that could see sky cues. In the same study, butterflies subjected to a 6 hour clock-shift treatment did not change their orientation. This is in contrast to migratory monarchs (Mouritsen and Frost, 2002) or hoverflies tested under similar conditions (Massy et al., 2021).

The red admiral, *Vanessa atalanta* (Linnaeus, 1758), is another Palaearctic butterfly species known to perform a fairly regular migration from the Mediterranean region toward the north in spring and summer and vice versa in autumn with a pronounced peak of butterflies being observed during autumn months according to ecological studies in different parts of Europe (Williams, 1951; Benvenuti et al., 1994; Stefanescu, 2001; Mikkola, 2003a; Brattström et al., 2008; Brattström et al., 2018). They usually overwinter in the Mediterranean region where they lay eggs on nettles (plants of the family Urticaceae) after arriving in autumn. However, hibernation of adults has not been reported there in winter (Stefanescu, 2001). According to previous recordings of vanishing bearings in the wild (Benvenuti et al., 1996; Stefanescu, 2001) and orientation behaviour in circular arenas (Brattström, 2007), we assume that this species can principally show appropriate migratory directions during spring, as well as autumn migrations. Yet, red admirals have never been tested in lab-controlled conditions that allow more in-depth investigations. Hence, despite their prominence, we know virtually nothing about the external compass cues that red admirals use as orientation references during their seasonal movements. For the first time in this study, therefore, we report an analysis of phenology data coupled with information about movement directions of more than 16,000 red admiral individuals recorded over 23 years of annual observations at the Baltic Sea. We further examined red admiral navigation behaviour during the autumn season, utilising both tethered butterfly recordings in a modified version of the Mouritsen-Frost flight simulator, and vanishing flight measurements gathered from translocation experiments that included a short, 5 km displacement from a coastal capture site. Generally, comprehensive, long-term studies of insect migration are scarce, yet they are of paramount importance. For example, only rigorous assessments of annual migration phenology enable the detection of shifts in mean migration counts which may be attributable to phenomena such as global warming (Cotton, 2003; Taylor, 2008). These shifts could potentially be associated with range shifts in butterflies (Parmesan et al., 1999; Sunde et al., 2023) which could, in turn, manifest as changes in orientation behaviours.

## Materials and Methods

### Phenology of butterfly movements

At the “Fringilla” field station (Kaliningrad region, Russia) red admirals and other butterfly species were recorded in “Rybachy-type” traps annually since 1982 from spring to autumn (between 1 April and 31 October, see details in Table S1). These large funnel traps passively catch aerial migratory animals, including insects (Brattström et al., 2018; Keišs et al., 2021). The field station operates two such traps, one facing to the northeast and the second to the southwest, so that the movement direction could be concluded for each red admiral individual. Using a northwest facing funnel trap at the Pape Ornithological Research Center (Latvia) large flying insects that could not escape the funnel trap were recorded between August 13th and October 31st, 2021, for the same purpose. Under favourable weather conditions, the trap was inspected in hourly intervals. After identifying the species and counting the individuals, we released the animals unharmed behind the trap. In addition to the red admiral, we focused on the mourning cloak (*Nymphalis antiopa*), the common brimstone (*Gonepteryx rhamni*), the large white butterfly (*Pieris brassicae*) and the European peacock (*Aglais io*) to describe the phenology of coastal butterfly movements. The temporal occurrence of five species of butterflies, including red admirals, is represented in Figure S6.

### Butterfly catching for experiments

Red admirals of both sexes were caught randomly in the wild during their autumn migrations, 2020-2021, using the traps described above, at “Fringilla” field station and the Pape Ornithological Research Center. The traps are operated daily from the beginning of April (“Fringilla”) and August (Pape), respectively, to the end of October, including the period when butterfly migration is ceasing at the Baltic Sea coast coinciding with the onset of the autumn storm season (see details in Brattström et al., 2018). Red admirals are non-Red Data book species according to the Federal Law on the Animal Kingdom (Russia) and in Latvia. Experimental procedures at “Fringilla” were approved by the Ethics Committee for Animal Research of the Scientific Council of the Zoological Institute, Russian Academy of Sciences (permit № 2020-7). The Pape trap was operated as part of the annual bird and bat migration research programmes.

While there are no official ethical requirements when working with insects at our field sites, we took great care when we handled butterflies. After experiments, and as part of experiments, all butterflies were released unharmed into the wild.

### Flight simulator experiments

#### Butterfly keeping

Upon capture, all butterflies were kept indoors (a laboratory chamber with windows) in mesh cages under natural photoperiods. Red admirals were fed with 10% honey solution (in water) and acclimatised for 3 days before tethering in the flight simulator.

#### A flight simulator and tethering

For condition-controlled experiments we used a modified version of a flight simulator constructed after (Mouritsen and Frost, 2002; Dreyer et al., 2021) to obtain butterfly flight tracks and analyse orientation. The flight simulator was made from non-magnetic materials (PVC) and is based on a white plastic cylinder (diameter 45 cm, height 50 cm; Fig 1) placed vertically on a plastic table. A miniature optical encoder E4T (US Digital, Vancouver, USA) was attached to a narrow rectangular aluminium tube and a cable connected the encoder and the data acquisition device or DAQ (USB4 model, US Digital, USA) was hidden inside this tube. The encoder recorded the butterfly’s heading (with a horizontal resolution of 1°) every 200 ms so we sampled 5 headings per second which were saved as .csv-file to an Asus PC tablet. A 15-cm long fine tungsten rod (diameter 0.5 mm) was connected to the encoder and created the encoder shaft, the end of which was attached to a butterfly during the trials. The edges of the white plastic cylinder restricted the butterflies’ visual access to the sky to 110°. We mounted a Raspberry Pi camera (miniature Pi v. 2.1 8MP 1080p camera module, connected to Raspberry Pi 4) in the centre of the plastic table to film the flight behaviour of the red admiral from below. The video from one of cameras was streamed to a Raspberry Pi 4 for real-time observation of butterfly’s flight behaviour and used as a backup source for analysis of orientation if USB4 software could not save data from the encoder. In 2021, we upgraded our setup: a 120-mm diameter 3D-printed plastic pipe with hundreds of 3 mm-holes was placed in the centre of the plastic table (instead of the Pi camera) to create laminar airflow from a computer fan (NF-F12, Noctua, Austria). This airflow can stimulate tethered butterflies to engage in active flight (Mouritsen and Frost, 2002). Two miniature video cameras (Waveshare Electronics, model H) were symmetrically placed around the encoder and recorded butterflies’ behaviour above. We used two cameras (instead of one) to create symmetrical environment inside the flight simulator.

**Figure 1.**
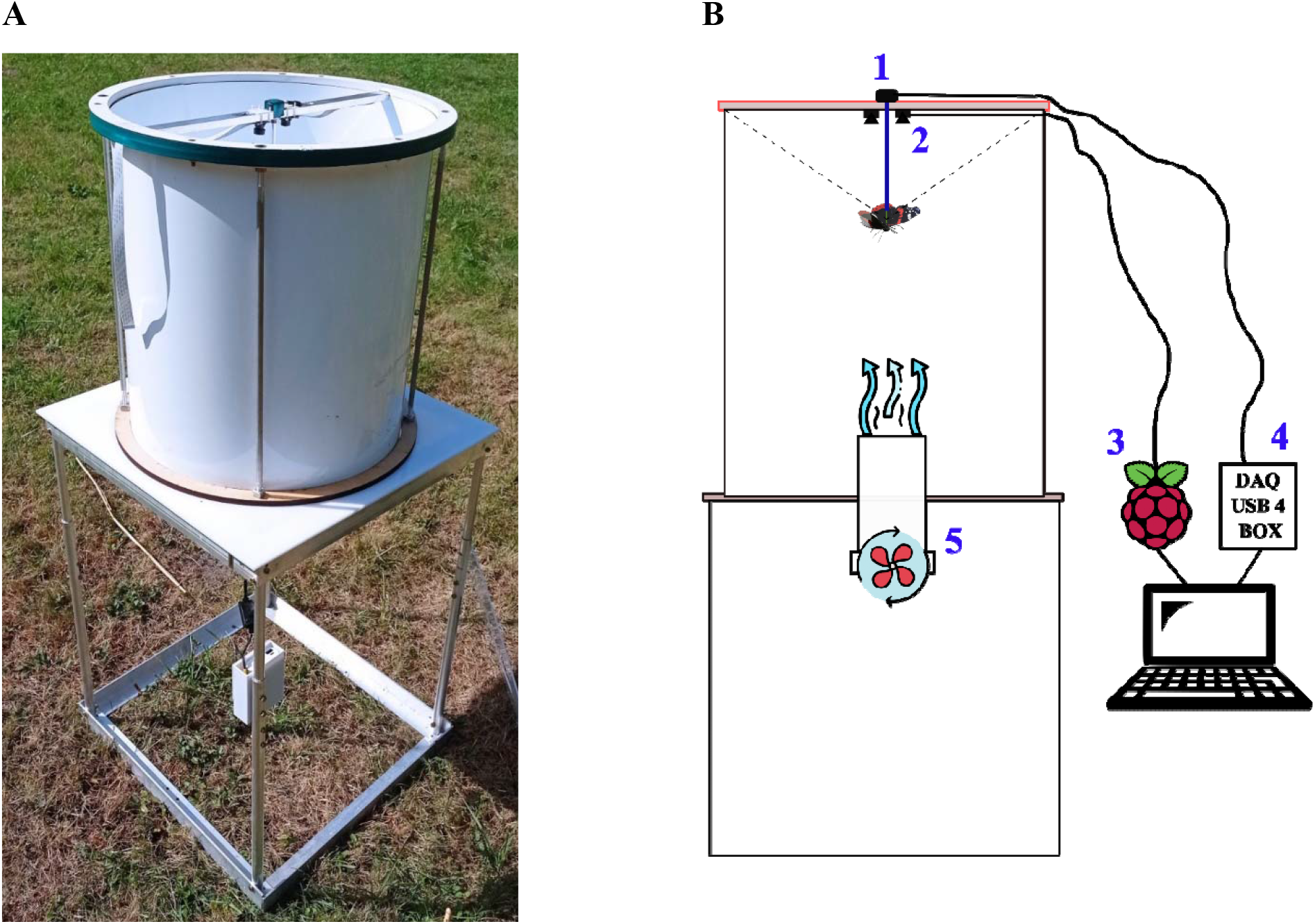
A photo (A) and scheme (B) of the modified version of Mouritsen-Frost flight simulator to study orientation and navigation in migratory butterflies. 1 -an optical encoder, 2 -a wide angle miniature camera, 3 -Raspberry Pi 4, 4 -data acquisition device (DAQ), 5 -a PWN computer fan.

All tethering procedures were non-invasive attachments (Parlin et al., 2021). Prior to attachment to the flight apparatus, each red admiral was kept in a fridge for 5-10 min to immobilise it. After removing of thorax’s scales, a small (3 cm) section of tungsten rod (ferromagnetic free) with footplate (attached to a small piece of paper: a tethering stalk) was vertically glued to this part of a butterfly using the ‘Bostik’ contact (Nesbit et al., 2009) or the non-toxic professional makeup Duo eyelash adhesive (Bojarinova et al., 2020). The butterflies were then returned to the mesh cage where they were kept at least one more day to recover from this procedure and before testing them in a flight simulator.

#### Orientation tests in-situ

All flights took place outside between 10:30 and 15:00 hours local time (GMT+2) in sunny condition (the sun’s disc was visible during all flight, without overcast). Prior to outdoor experiments, we placed mesh cages with tested butterflies outside on the experimental site in direct sunlight for at least 30 min before testing. This procedure allowed them to warm up and sense different orientation and navigation cues before testing. Each red admiral was taken out of the mesh cage by grasping the stalk attached to the thorax. This stalk was then inserted into a small plastic tube (20 mm length, diameter 2 mm, 0.5 mm hole) attached to the encoder shaft. After attaching to the encoder shaft, each red admiral individual was first aligned by hand to magnetic North (mN) and the encoder was reset to zero via USB4 software. Each test lasted 15 min and was included in further analysis only if a butterfly flew 5 min at least. We omitted the first minute of each flight to avoid any handling bias on orientation of insects. All experiments were carried out in rural locations with a low level of radio-frequency noise (Pakhomov et al., 2017) and our experimental setup did not produce radio-frequency disturbances at the levels that would theoretically disrupt ability of tested specimens to orient in appropriate migratory direction (Figure S1). All measurements were performed using a calibrated active loop antenna FMZB 1512 (Schwarzbeck Mess-Elektronik, Germany) and a spectrum analyzer Rigol DSA815-TG (Rigol Technologies, USA).

### Release experiments

#### Butterfly keeping

Upon capture, red admirals obtained using the trap at the Pape Ornithological Research Center in 2021 were kept indoors overnight, in mesh cages. Acclimatisation lasted at least until noon the following day, i.e., until transportation to the release site. Under bad weather conditions that would not have allowed for releases because of wind and or rain, butterflies in captivity became torpid. On such days, we could observe that the migration of wild red admirals came to a halt. The keeping was then prolonged until wild butterflies were seen migrating again, i.e., when the first experimental release was possible again also. As a consequence of experimental releases, butterflies were free, back in the wild.

#### Translocation, release, and orientation recording under free-flight conditions

Red admirals were displaced to a forest meadow 5 km east of the coastline (Lindecke et al., 2019). This meadow was chosen because it offers a radius of 70 m uniform habitat in all directions (grass cover), i.e., without any higher vegetation that would be neither obstructing the view for the animals nor providing landmarks to them in close range. We released butterflies between 13:00 to 14:00 (21-24 August, 2021), around the time of local solar noon, i.e., when the Sun passed the meridian and reached the highest position in the sky (α_AZ_ = 177°, NOAA Solar Calculator). Each butterfly was carefully taken out of the fabric mesh holding cage and released one at a time from 2 m height above ground. Red admirals could take off in their own time and under wind still to 1 m/s wind conditions with clear sky and or the occasional cloud. Butterflies were visually tracked until they vanished from view at the position of release, i.e., at least for a 60-70 m radius. Vanishing bearings were recorded using a handheld compass.

### Data acquisition and statistical analysis

Only butterflies flying actively in a flight simulator for at least 5 min during each 15 min test were included in the data analysis. For each individual flight obtained by an optical encoder in 2020, we calculated the mean flight direction (α) of the individual red admiral and the directedness of its flight, individual r value (mean orientation vector length). In 2021, owing to the problems with saving data collected by an optical encoder, we used video files as a primary source to analyze direction of flights. We performed a video analysis using Python with additional libraries: *opencv, numpy, matplotlib, scikit-learn, pytorch*. The whole model for the video analysis (Fig. S2) aims to answer two key questions: when was a butterfly active and how did it orient itself during these periods of activity. In order to analyze butterfly orientation, two machine learning models were used. The first artificial neural network (ANN1) was based on ResNet18 architecture with modified fully-connected layers. For the second model (ANN2) a simple straight-forward network with a single convolution and three fully-connected layers was created (Table S3). Both nets are intended to perform a classification task, ANN1 has 8 output classes (Fig. S3; Table S4), and ANN2 is a binary classification model (accuracy 0.98, f1-score: 0.967).

There is no consensus among researchers on the threshold r value for flight path directionality, whether it is r > 0.1 (Mouritsen and Frost, 2002) or r ≥ 0.2 (Nesbit et al., 2009; Dreyer et al., 2018). Therefore, we have decided to use all data with r > 0.1. Other criteria of directedness, such as the Z-score (Z = nr^2^, where n is the number of observations and r is the magnitude of the mean vector (Zhu et al., 2009)), are not applicable to our data since we have Z-scores only for data collected by the optical encoder and the DAQ USB4 device in 2020. The group mean vector was calculated by vector addition of unit vectors in each of the individual butterfly’ mean directions. The classical Rayleigh test (RT) was used to compare the group mean orientation against uniformity (Batschelet, 1981). Additionally, we were able to conduct second order statistics for this purpose -the nonparametric Moore’s modified Rayleigh test (MMRT; (Moore, 1980) which allows to weight the mean angles according to their r value. For both tests (RT and MMRT) low p-value (p < 0.05) indicates the tested butterflies chose preferred direction. The data of the release experiments was analysed using the RT, as well. Data analysis was conducted using Oriana 4.01 (Kovach Computing Services, UK).

## Results

### Orientation of red admirals during autumn migration

Red admirals in the flight simulator were oriented in a southwestern direction under the clear sunny sky and with access to the undisturbed geomagnetic field (α = 198°, r = 0.498, n = 19, 95% CI = 164°–232°, RT: Z = 4.7, p = 0.007, Fig 2; second order statistics: α = 210°, r = 0.26, n = 19, 95% CI = 145°–264°, MMRT: R^*^ = 1.136, p < 0.025, Fig. 3A). If we included only individual mean directions with r ≥ 0.2 in the evaluation, which would indicate more highly directed flights (Nesbit et al., 2009; Dreyer et al., 2018), the results do not change: butterflies continue to show a seasonally appropriate migratory direction (α = 197°, r = 0.455, n = 17, 95% CI = 157°–237°, RT: Z = 3.5, p = 0.027, Fig 3B; second order statistics: α = 210°, r = 0.28, n = 17, 95% CI = 141°–267°, MMRT: R^*^ = 1.1, p < 0.05, Fig. 3C) and no statistically significant difference is found between these directions (MWW test: W = 0.077, p = 0.96).

**Figure 2.**
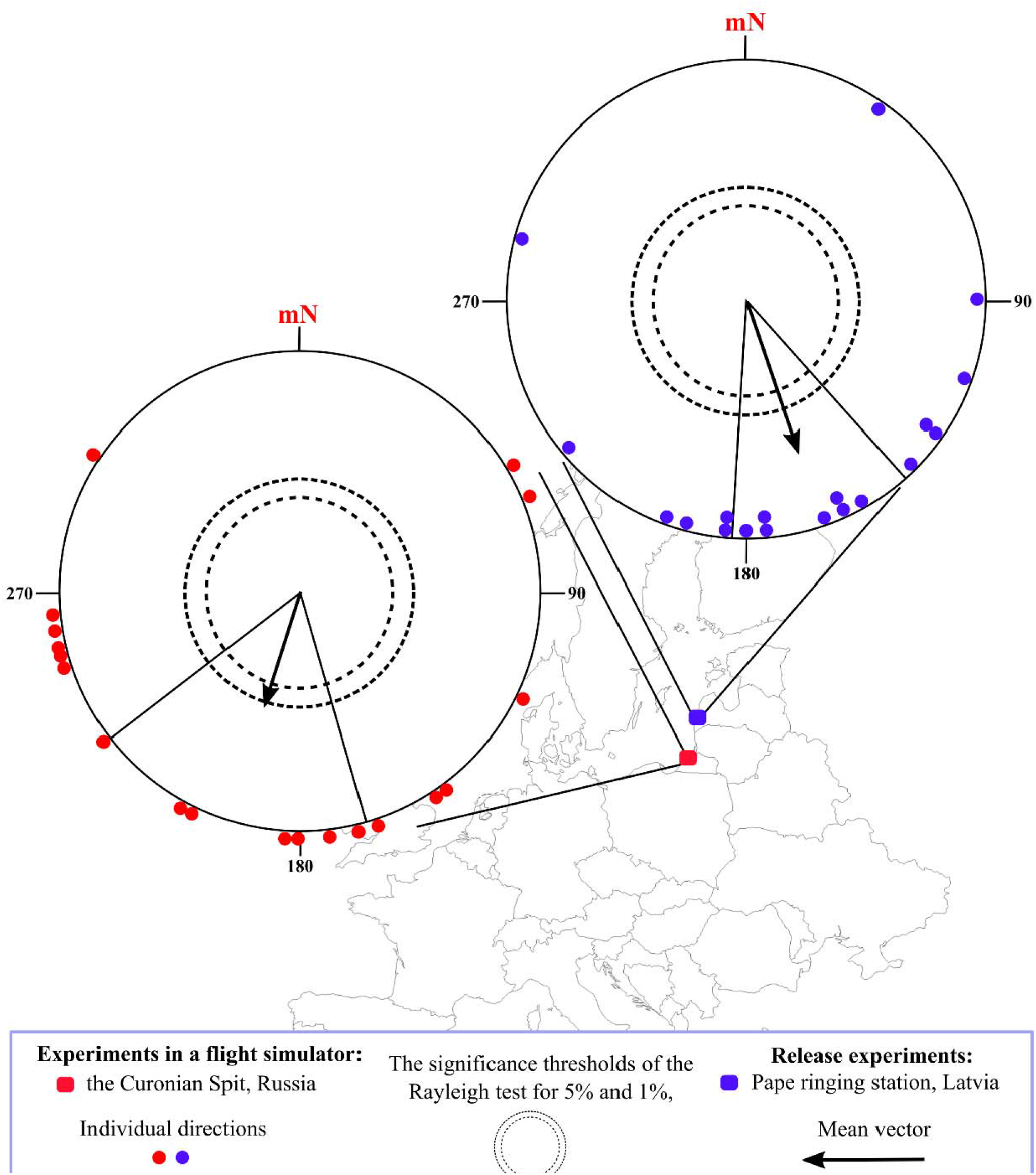
Orientation of red admirals from the Baltic region observed in a flight simulator and in free-flight after translocation and release. Dashed lines indicate the significance thresholds of the Rayleigh test for 5% and 1%, respectively. Each dot at the circle periphery indicates the mean orientation of one individual butterfly.

**Figure 3.**
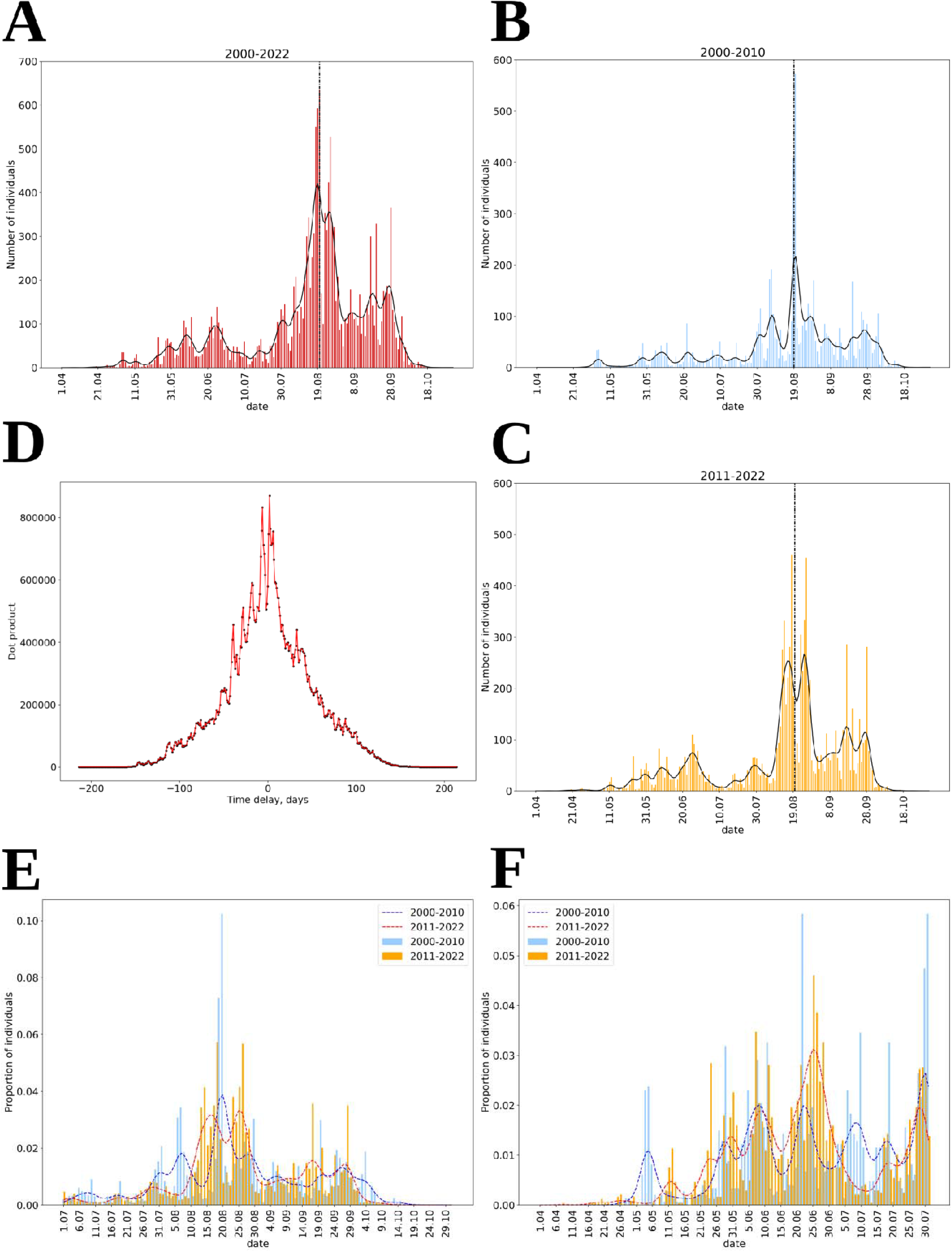
Seasonal abundance of red admirals captured by the stationary trap at the Fringilla field station, the Courish Spit (Russia). **A)** The number of red admirals captured per day in the “Rybachy-type” traps for 23 complete seasons (April-October) 2000-2022. **B)** and **C)** Graphs showing the same data with two time-periods (2000-2010 and 2011-2022) plotted separately with median dates of capture (dashed lines). **D**) Cross-correlation plot for the number of individuals per day for 2000-2010 and 2011-2022 periods; **E**) Proportion of individuals per day relative to the total number of individuals caught in April-July for 2000-2010 period (blue bars, the blue dotted line represents the global trend) and 2011-2022 period (orange bars, the red dotted line represents the global trend); **F**) Proportion of individuals per day relative to the total number of individuals caught in July-October for 2000-2010 period (blue bars, the blue dotted line represents the global trend) and 2011-2022 period (orange bars, the red dotted line represents the global trend).

Red admirals released in free-flight trials were oriented in a south-southeastern direction (α = 161°, r = 0.68, n = 19, 95% CI = 138°–183°, RT: Z = 8.8, p << 0.001, Fig. 2). This direction runs parallel to the local coastal migration flyway; however, the butterflies due to the translocation had no access to coastal landmark cues.

### Phenology of the movements of red admiral

In total, 20,432 individuals of red admiral were recorded in 41 years of annual observations conducted at the “Fringilla” field station in 1982-2022 (a detailed data on recorded specimens and associated parameters, such as wind direction, cloudiness, air temperature is given in the Table S5). The highest number of red admirals was recorded in the year 2022 (a total of 2,705 captured individuals). In further analysis, we restrict ourselves to 16,133 records of red admiral gained in 2000-2022, when full-season observations (from the beginning of April to the end of October) were conducted. The estimated median date of red admirals’ passage over the Courish Spit is 20 August (Figure 3A). In order to investigate migration patterns between years and to reveal possible variation in the migration dates over the time, we estimate and compare median capture dates for two following periods: 2000-2010 and 2011-2022. Both in 2000-2010 (period 1) and 2011-2022 (period 2), the peak of migration was observed in late summer (median date for 2000-2010: 20.08; median date for 2011-2022: 19.08; Figures 3B,C). Cross-correlation analysis suggests that patterns of butterfly migration for period 1 and period 2 are best fitted to each other with no time delay (Figure 3D). The most noticeable difference between both periods is the appearance of an increase in the number of individuals in the early summer of period 2 (Figure 3E). We did not observe any other apparent differences in autumn migration patterns (Figure 3F). The revealed median date also suggests that we conducted our experiments on navigation behaviour when migration movements generally peak at our study sites. In order to investigate seasonally preferred movement directions of red admirals over the Courish Spit, we limited analyzing records to dates when two “Rybachy-type” traps of opposite directions (facing to the northeast and to the southwest) were operated. Varying captures of red admirals are evidence for generally movement pattern of the species changes drastically over the season. While in April, May, and June butterflies flew in northeast and south west directions in nearly equal proportions, in August, September, and October red admirals strongly favoured southeast direction (Figure 4).

**Figure 4.**
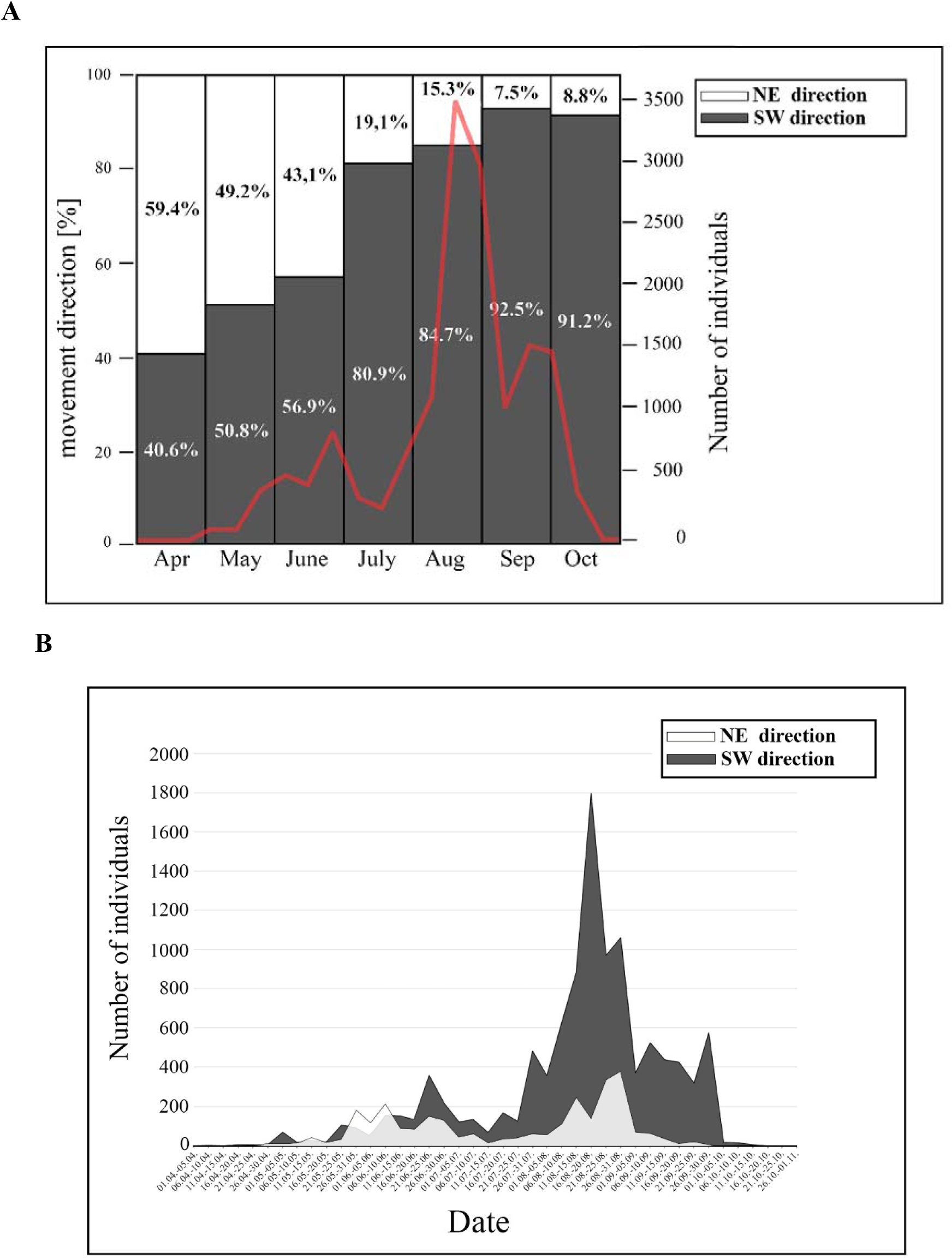
Directional preferences of movements of red admirals (A -proportions, B – number of individuals) captured in two large traps (see Figure S1 for details). Light green colour – NE movements, % of red admirals caught by the NE trap; purple colour – SW movements, % of red admirals caught by the SW trap. The red line depicts the number of caught butterflies based on weekly count sums.

## Discussion

The characterization of long-range movement behavior of the red admiral has undergone significant shifts over the years: it was categorized as a resident species at the dawn of the 20th century, redefined as a short-distance migrant by the century’s midpoint (Williams, 1951), and ultimately identified as a long-distance migratory species in the early stages of the 21st century (Mikkola, 2003a). Every autumn they migrate to the Mediterranean region where they lay eggs on nettles. When the new generation of red admiral emerges in spring, they migrate northward and then lay eggs again on the host plant. One or two further summer generations of red admirals continue to recolonise Northern Europe thereafter (Stefanescu, 2001), and in autumn a main mass of red admirals starts their southward migration towards the wintering places in Southern Europe and North Africa. Due to this full annual cycle, two peaks of abundance of red admirals can be recorded in different parts of Europe and the number of butterflies in autumn is actually dramatically larger than in spring (Benvenuti et al., 1994; Stefanescu, 2001; Brattström et al., 2018). Our long-term recordings of migrating red admirals on the Courish Spit correspond with this pattern: first butterflies appear here in the second half of April and a first small peak of adults is seen in June (Figure 3). After that, the numbers decrease and remain low until the second larger peak in the middle of August.

In the experimental parts of our study, red admirals on the Courish Spit of the Baltic Sea were observed to orientate in a southwestern direction in flight simulators. One hundred kilometres further up the coast in Latvia, red admirals vanished in a south-southeastern direction when released in free-flight trials after translocation from the dunes of the Baltic Sea. Data from previous studies documenting the vanishing directions observed in freely moving butterflies indicate that red admirals typically choose a certain range of directions from east to southwest across different parts of Europe. These directions include south preferences in the British Isles, Denmark, and Finland (Williams, 1951; Hansen, 2001; Mikkola, 2003b); southwest preferences in Germany, Sussex, Spain, and Sweden (Roer, 1991; Leverton, 2000; Stefanescu, 2001; Brattström, 2007; Brattström et al., 2008); and southeast and east preferences in Italy (Benvenuti et al., 1994; Benvenuti et al., 1996). Hence, our results match the general autumnal migration directions in Europe, suggesting that both the flight simulator approach and translocation trials, as well, qualify as tools for investigating navigation behaviors in red admirals.

Probably, in contrast to monarch butterfly in North America, red admirals do not have certain winter places, do not form aggregations and usually terminate their migration when they reach a suitable place for mating and laying eggs on host plants (Mikkola, 2003a). Red admiral is a typical daytime FBL migrant whose migration take place close to the ground within so-called “flight boundary layer” or FBL (Chapman et al., 2015; Chowdhury et al., 2021) with rare observations of high-altitude migration (Mikkola, 2003b). Flying within FBL has several advantages for insects: they can avoid predators, initiate their migration whenever environmental conditions allow (optimal temperature, access to orientation cues, etc.) and control preferred fly direction without influence of prevailing wind direction (Brattström et al., 2008; Chapman et al., 2015). This butterfly species seem to select an adaptive migratory direction following coastlines and trying to avoid large bodies of water (Brattström et al., 2008) and this idea can explain difference in direction between relatively close study site such as Falstebro in Sweden (SW and W directions) and Denmark (S direction) (Hansen, 2001; Brattström, 2007; Brattström et al., 2008), two orientation sites in central-northern Italy (SE and E directions) (Benvenuti et al., 1994; Benvenuti et al., 1996) and our study sites too. Red admirals as other diurnal migrating lepidopterans (Schmidt-Koenig, 1985; Vander Zanden et al., 2018) may rely on information from the Sun and/or polarized light as primary orientation cues (Perez et al., 1997; Mouritsen and Frost, 2002; Nesbit et al., 2009) but frequently use the sea coast as a guiding line. Therefore, our butterflies in the Pape Nature Park flew in southeastern direction along the SE coast even after short-distance physical displacement and demonstrated SW orientation (along the NE-SW axis of the Courish Spit) when they were tested in the flight simulator. Similar to results of studies in migratory birds tested in Emlen funnels (Nievergelt et al., 1999; Bäckman and Alerstam, 2003), wild caught red admirals did not show a highly concentrated orientation in our flight simulator experiments in contrast to release tests, probably, due to unnatural test condition and experimental manipulation (tethering, testing inside the simulator without any visual landmark cues, etc.) or reverse migration of some testing butterflies. Reverse migration in other migratory animals, for example, birds, usually occur before the ecological barriers (open sea water, deserts, etc.) if they have low fat reserve to cross these barriers successfully (Sandberg, 1994; Åkesson et al., 1996). As can be seen in Figure 4, some kind of reverse migration of red admirals in autumn occurs on the Courish Spit, too: in spring and summer red admirals fly in northeast and southwest directions in nearly equal proportions, but during autumn migration most of them prefer southwestern direction but 7-15 % of all butterflies are captured by our NE trap. In our flight simulator experiments we used only butterflies caught in the SW trap but we cannot exclude that some red admirals flying in northern direction can be collected by the NE trap too. Additionally, there is evidence that this species can fly northward in circular cages (Brattström, 2007) and the scattered orientation has been shown on other migratory butterflies and moths in flight simulator experiments (Nesbit et al., 2009; Dreyer et al., 2018).

Regular annual migrations, large numbers of butterflies in autumn every year, and proper orientation under lab-controlled conditions make red admirals a good model species to study orientation and navigation in European butterflies. However, there is a lack of our knowledge about compass and navigation systems in Palaearctic migratory butterflies: we can only say that they might use the sun compass to choose and maintain appropriate migratory direction (Brattström, 2007; Nesbit et al., 2009). This contrasts with the fact that butterflies in Europe are popular models in a broad range of ecological studies because they are commercially important pollinators besides honey bees, wasps, hoverflies, and bumblebees. Reproductive success of many wild and economically important plants depends on pollination by animals (Potts et al., 2016). Despite their value, populations of pollinators, especially insects, have declined dramatically in North America and parts of Europe (Koh et al., 2016; Powney et al., 2019). Currently, both two most iconic model species for studying orientation and navigation in Lepidoptera, North American monarch butterfly and Australian Bogong moth *Agrotis infusa* (Warrant et al., 2016; Reppert and de Roode, 2018), are in global endangered species list: monarch butterfly population has declined over the last two decades (Thogmartin et al., 2017) and population of Australian moths has crashed rapidly in recent years. Red admirals are not in IUCN Red List of Threatened Species yet and the numbers of butterflies migrating in the Baltic region do not decrease year after year during the last two decades according to the results of our study, but this situation can change in the future. Our knowledge about how red admirals from Northern Europe find a way to the Mediterranean region and vice versa, what factors initiate or terminate their southward and northward migrations and how their compass systems work help us to understand how climate change and environmental pollution (artificial light at night, electromagnetic noise, etc.) caused by human activity can have impact on life of these tiny creatures.

## Supporting information

Supplementary materials (pictures)

## Acknowledgments

We are grateful to Ivo Dinsbergs and Dr. Oskars Keišs (Institute of Biology, University of Latvia) for supporting our fieldwork at Pape station, to Roman Cherbunin for his help in measurements of radio-frequency noise.

## Authors’ contributions

A.P. designed flight simulator research; A.P. performed experiments in a flight simulator, collected and analyzed the data; F.C. wrote Python code for neural network to analyze video; O.L. and R.J. conceived and performed release experiments; N.S. and A.S. collected butterflies in Kaliningrad region and analyzed phenology data, whereas R.J. and O.L. did the same for butterflies in Pape, Latvia; A.P., N.S. and O.L. wrote the first draft of the manuscript. All authors commented on the manuscript and gave final approval for publication.

## Funding

Financial support for this study was made available by the Russian Science Foundation (grant 21-74-00093 to A.P., flight simulator experiments in 2021) and Zoological Institute RAS (research projects 12203110026-7 to A.P. and 122031100272-3 to N.S.). Fieldwork by O.L. was partially supported through the Sonderforschungsbereich (SFB) 1372 (project-ID 395940726) by the Deutsche Forschungsgemeinschaft (DFG). O.L and R.J. further received funding from the Baltic-German University Liaison Office (grant 7/5-2022).

## Competing interests

The authors declare no competing interests.

## Data Availability

All data generated or analyzed during this study are included in this manuscript (and its Supplementary Information files) or available from the corresponding author on reasonable request.

## Notes

### Competing Interest Statement

The authors have declared no competing interest.

