## Supplementary materials (pictures) for "Orientation tests and long-term movement phenology establish the red admiral *Vanessa atalanta* as an applicable model for navigation research in migratory butterflies"

**Figure S1. Photograph of the large ‘Rybachy-like’ traps (the Courish Spit, Russia) from a drone (from different positions) which are used to passively catch migratory birds and insects. Arrows indicate direction of each trap, 0˚ is the magnetic North.**

NE trap: for birds and butterflies which fly in northern direction, SW: for birds and butterflies which fly in southern direction.

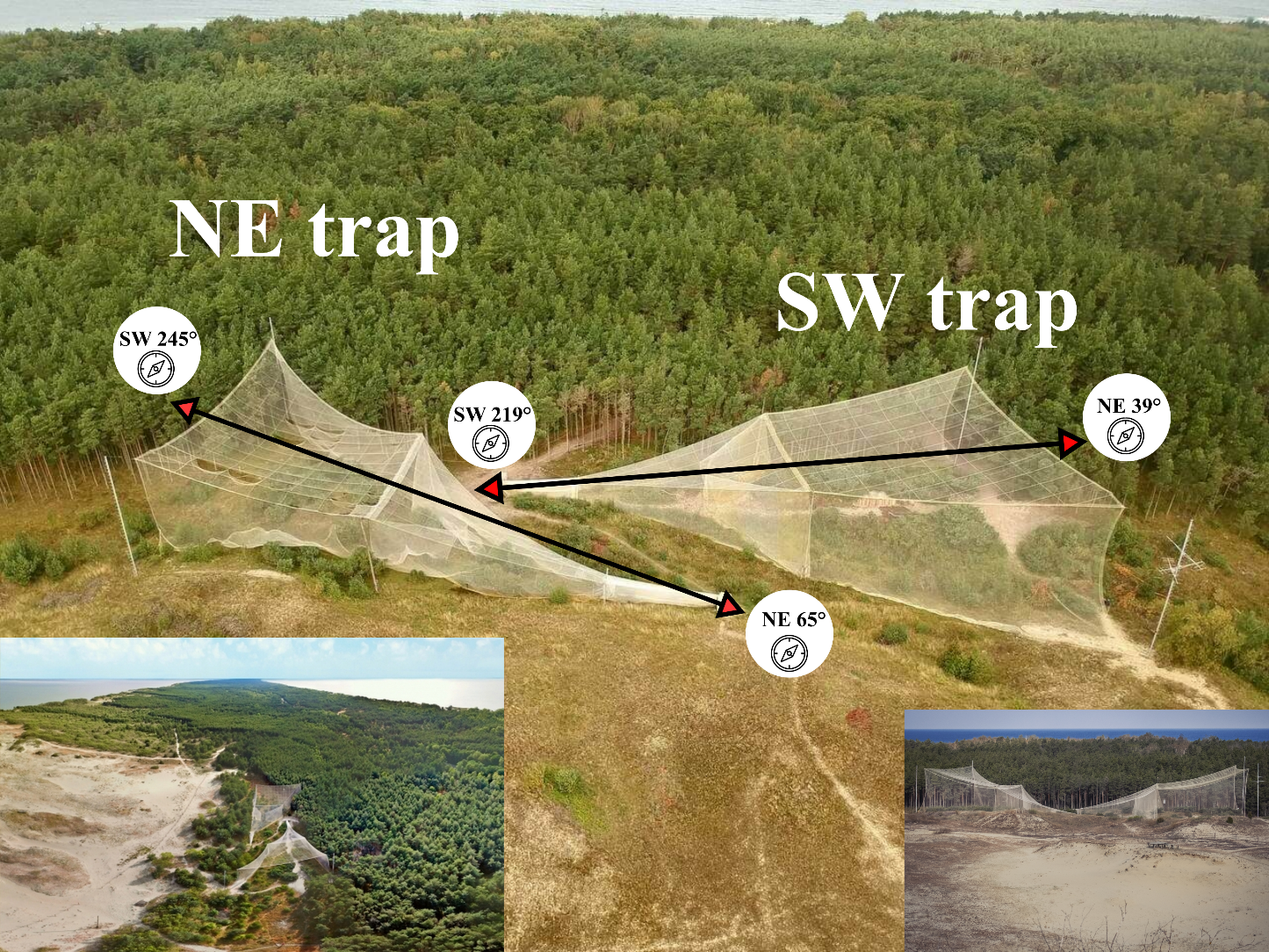

**Figure S2. Average amplitude of the magnetic field noise measured at the experimental site as a function of the central position of the 10 kHz detection window.**

1 – near the box with minicomputer Raspberry PI (2 m away from the flight simulator)

2 - inside the flight simulator, the place where red admiral was attached to a vertical encoder shaft; all electronic equipment was turn on

3 - inside the flight simulator, the place where red admiral was attached to a vertical encoder shaft; all electronic equipment was turn of

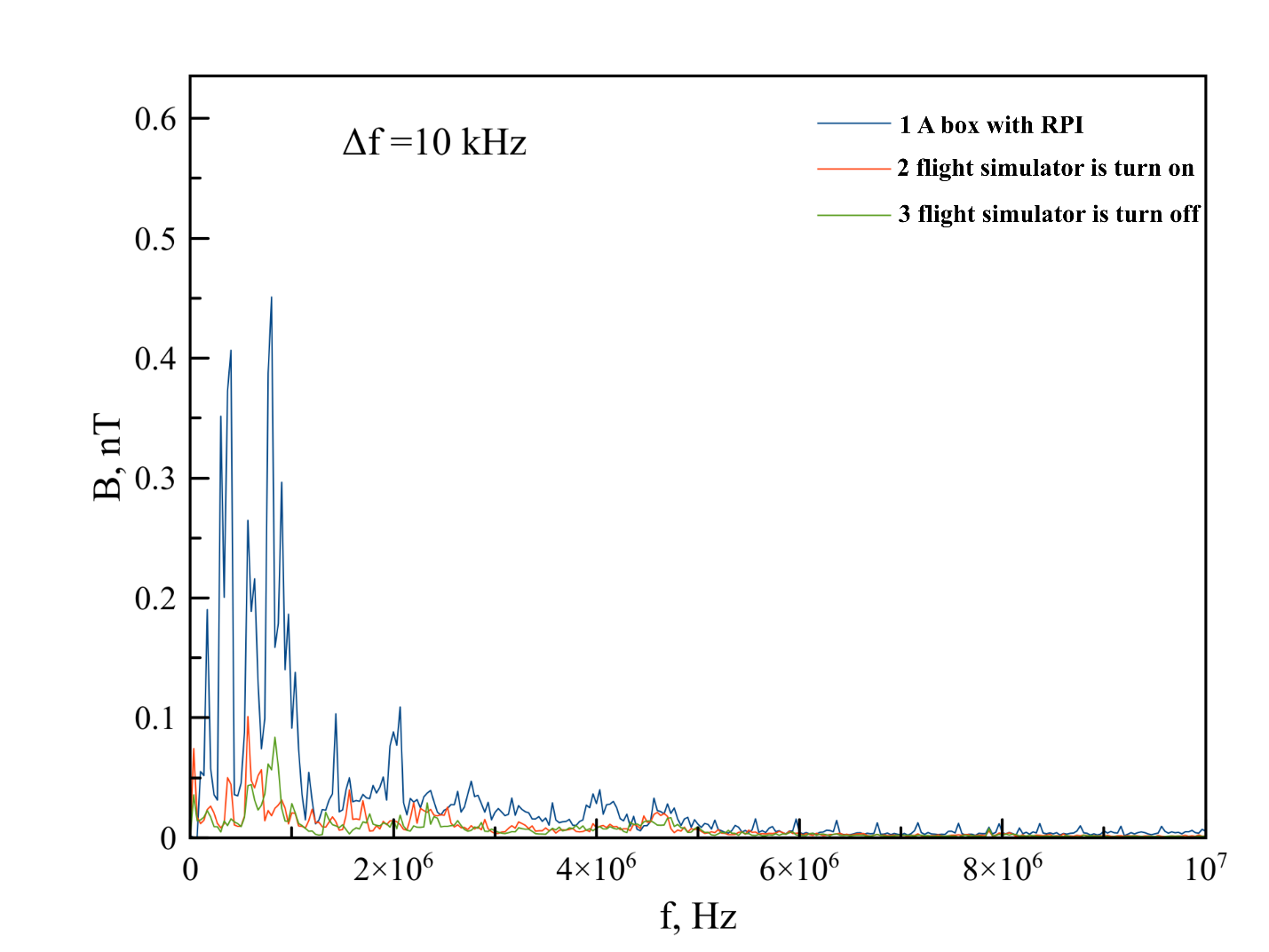

**Figure S3. The overview of a neural network-based model to analyze behaviour of butterflies tested in a flight simulator.**

The whole video analysis aims to answer two key questions: when a butterfly was active and how did it orient during this activity periods. At the beginning of the analysis, an original video (**a**) was cropped roughly to the size of a butterfly (**b**). As a butterfly consequently opens and closes wings during the active flight, frame brightness changes respectively. The means for each frame by the three RGB channels clearly show when the butterfly implies active flight (plot **c**: frame number by the X-axis, means by channels by Y-axis). In order to define for each frame, is a butterfly flying on it or not, a 50-frame window (starting with the frame of interest) was selected and the range of means was calculated for the window. Values of the range were divided into two classes by the K-means algorithm. These classes correspond to active flight and periods of inactivity (plot **d**: frame number by the X-axis; the upper line represents the ranges, and the lower line represents the classes, distinguished by K-means).

Only frames where the butterfly was assumed active were used in further analysis. Each of these frames was classified by an artificial neural network (**ANN2**) to select ones where butterfly wings were opened. Such frames were further divided by another neural network (**ANN1**) into 8 classes, which corresponded to eight sectors on the circle. In the end, a track for each butterfly was created based on ANN1 labels (**e**).

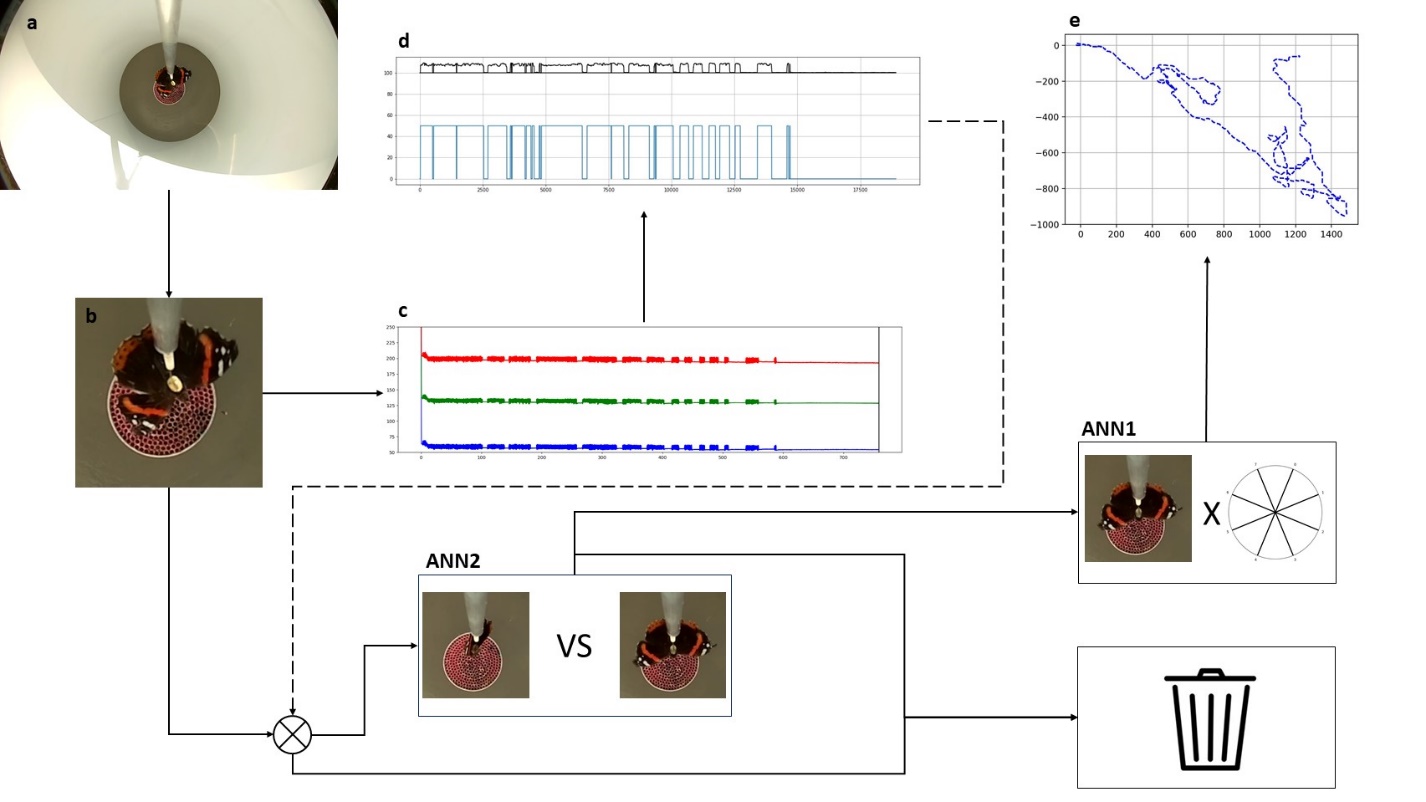

**Figure S4**. Graphical representation for ANN1 confusion matrix (rows represent the actual labels, columns predicted ones).

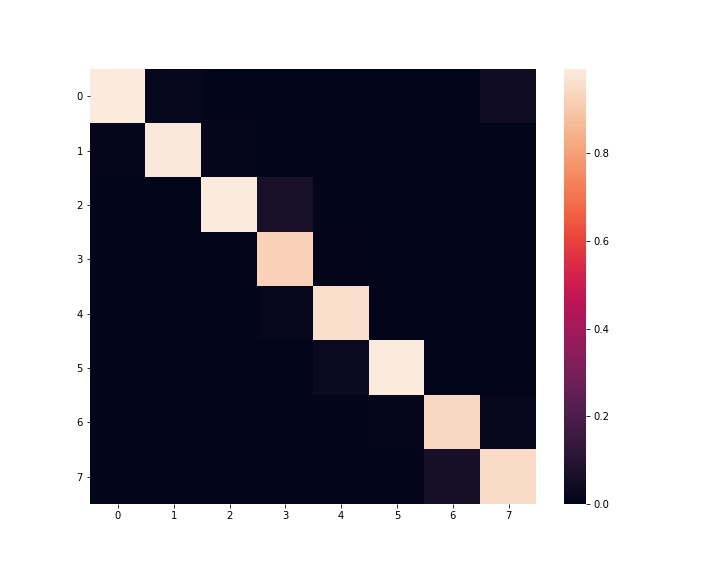

**Figure S5. The orientation of red admirals in the flight simulator experiments.**

**A – a circular diagram based on results of Moore’s modified Rayleigh test (all data), B - a circular diagram based on results of Moore’s modified Rayleigh test (r ≥ 0.2), C - a circular diagram based on results of the classical Rayleigh test (r ≥ 0.2).**

| **A**  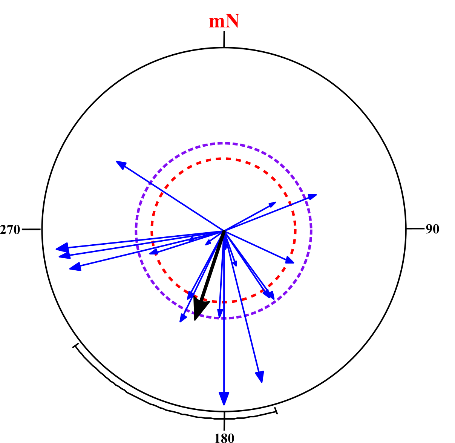 | **B**  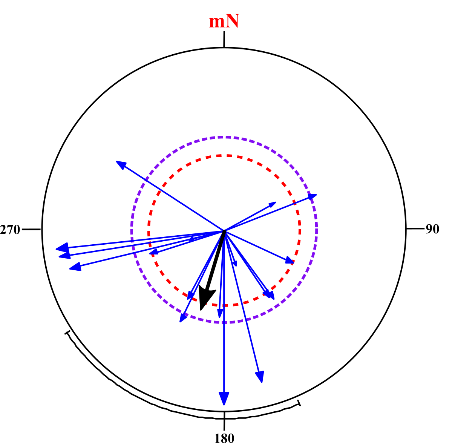 | **C**  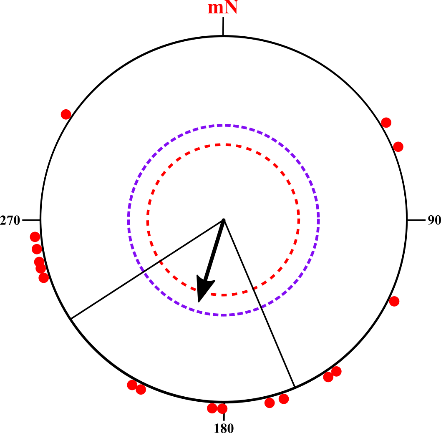 |
| --- | --- | --- |

**Figure S6. Abundance of several butterfly species captured by the stationary trap at the Pape Ornithological Research Center, Latvia, in 2021.**

No. of individuals

| Date (day.month) in 2021 | Vanessa atalanta | Nymphalis antiopa | Gonepteryx rhamni | Aglais io | Pieris brassicae | All species |
| --- | --- | --- | --- | --- | --- | --- |
| 2021-08-14 |  | 1 |  |  |  | 1 |
| 2021-08-22 | 7 | 1 | 6 |  |  | 14 |
| 2021-08-23 | 8 |  | 10 |  |  | 18 |
| 2021-08-27 | 13 |  |  |  |  | 13 |
| 2021-08-28 | 18 |  |  |  |  | 18 |
| 2021-08-29 | 19 |  | 26 |  |  | 45 |
| 2021-08-30 | 12 | 1 | 4 |  |  | 17 |
| 2021-08-31 | 7 | 2 |  |  |  | 9 |
| 2021-09-01 | 1 |  |  |  | 7 | 8 |
| 2021-09-03 |  |  | 2 |  |  | 2 |
| 2021-09-04 |  |  | 2 |  | 1 | 3 |
| 2021-09-05 |  |  | 2 |  |  | 2 |
| 2021-09-07 |  | 2 | 10 | 9 |  | 21 |
| 2021-09-08 |  |  |  | 2 |  | 2 |
| 2021-09-09 |  | 1 | 2 |  |  | 3 |
| 2021-09-10 |  |  |  | 7 | 4 | 11 |
| 2021-09-26 |  | 5 | 3 | 15 |  | 23 |
| 2021-09-27 |  |  |  | 1 |  | 1 |
| 2021-10-02 |  | 1 |  | 1 |  | 2 |
| 2021-10-03 |  |  |  | 2 |  | 2 |
| 2021-10-05 |  | 3 |  | 1 |  | 4 |
| 2021-10-09 |  |  |  | 1 |  | 1 |
| 2021-10-10 |  |  |  |  | 2 | 2 |

**Table S3. Artificial neural networks short descriptions.**

| model | description | n classes | input | training dataset size (balanced) | test dataset size | accuracy | balanced accuracy |
| --- | --- | --- | --- | --- | --- | --- | --- |
| ANN1 | Pretrained ResNet18 with modified fully connected layers | 8 | 224x224 3-channel image | 1056 | 1056 | -- | 0.97 |
| ANN2 | Simple straight-forward model with single convolution layer and 3 fully-connected layers | 2 | 64x64 3-channel image | 1500 | 376 | 0.98 | -- |

**Table S4.** ANN1 confusion matrix**.**

|  | Predicted | **0** | **1** | **2** | **3** | **4** | **5** | **6** | **7** |
| --- | --- | --- | --- | --- | --- | --- | --- | --- | --- |
| Actual |  |  |  |  |  |  |  |  |  |
|  | **0** | 125 | 2 | 0 | 0 | 0 | 0 | 0 | 5 |
|  | **1** | 1 | 130 | 1 | 0 | 0 | 0 | 0 | 0 |
|  | **2** | 0 | 0 | 123 | 9 | 0 | 0 | 0 | 0 |
|  | **3** | 0 | 0 | 0 | 131 | 1 | 0 | 0 | 0 |
|  | **4** | 0 | 0 | 0 | 2 | 130 | 0 | 0 | 0 |
|  | **5** | 0 | 0 | 0 | 0 | 4 | 128 | 0 | 0 |
|  | **6** | 0 | 0 | 0 | 0 | 0 | 1 | 129 | 2 |
|  | **7** | 0 | 0 | 0 | 0 | 0 | 0 | 8 | 124 |
